# Decline in mitochondrial health and lysosomal acidification in human neurons during normal aging

**DOI:** 10.64898/2026.04.09.717473

**Authors:** Emilie M. Legault, Juan-Pablo Izabal Gamez, Janelle Drouin-Ouellet

**Affiliations:** Faculty of Pharmacy, University of Montreal, Montreal, Quebec, Canada; Center for Interdisciplinary Research on Brain and Learning (CIRCA), Montreal, Quebec, Canada

**Keywords:** direct neuronal reprogramming, induced neurons, mitophagy, aging, mitochondria, mitochondrial health, human neurons

## Abstract

**Introduction:** In humans, aging is associated with an increased risk of developing neurodegenerative diseases such as Parkinson’s disease (PD) and Alzheimer’s disease (AD). In neurons, the effect of aging on intrinsic molecular processes, and how they tie to age-related neurodegeneration remains unclear. Animal studies have shown that mitochondrial function decline, autophagy impairment and defective elimination of damaged mitochondria by mitophagy are all central features of neuronal aging. However, very few studies have investigated such events in human neurons, due to a lack of models showing aging features. Therefore, there is a crucial need for a better understanding of the effect of aging on neuronal health.

**Methods:** Here, we use direct neuronal reprogramming, which maintains signatures of cellular aging, to study the effect of aging on mitochondrial health and mitophagy in human neurons derived from healthy donors ranging from 11 to 91 years old. We use confocal imaging to assess colocalization of mitochondria with autophagic structures following mitophagy induction.

**Results:** We show age-related mitochondrial impairment in neurites of induced neurons (iNs) derived from older donors. Additionally, in these neurites, we show an accumulation of mitochondria targeted for degradation in autophagosomes and LAMP1-positive lysosome vesicles following mitophagy induction. These impairments culminate into incomplete elimination of damaged mitochondria, despite preserved activity of the lysosomal enzymes assessed.

**Conclusions:** This study shows for the first time age-related mitophagy impairment in human neurons, and confirms findings of age-related mitochondrial impairments in human neurons. By doing so, this study paves the way for more in-depth mechanistic studies that would allow for the identification of therapeutic targets for anti-aging treatment and in the context of age-associated neurodegenerative diseases.

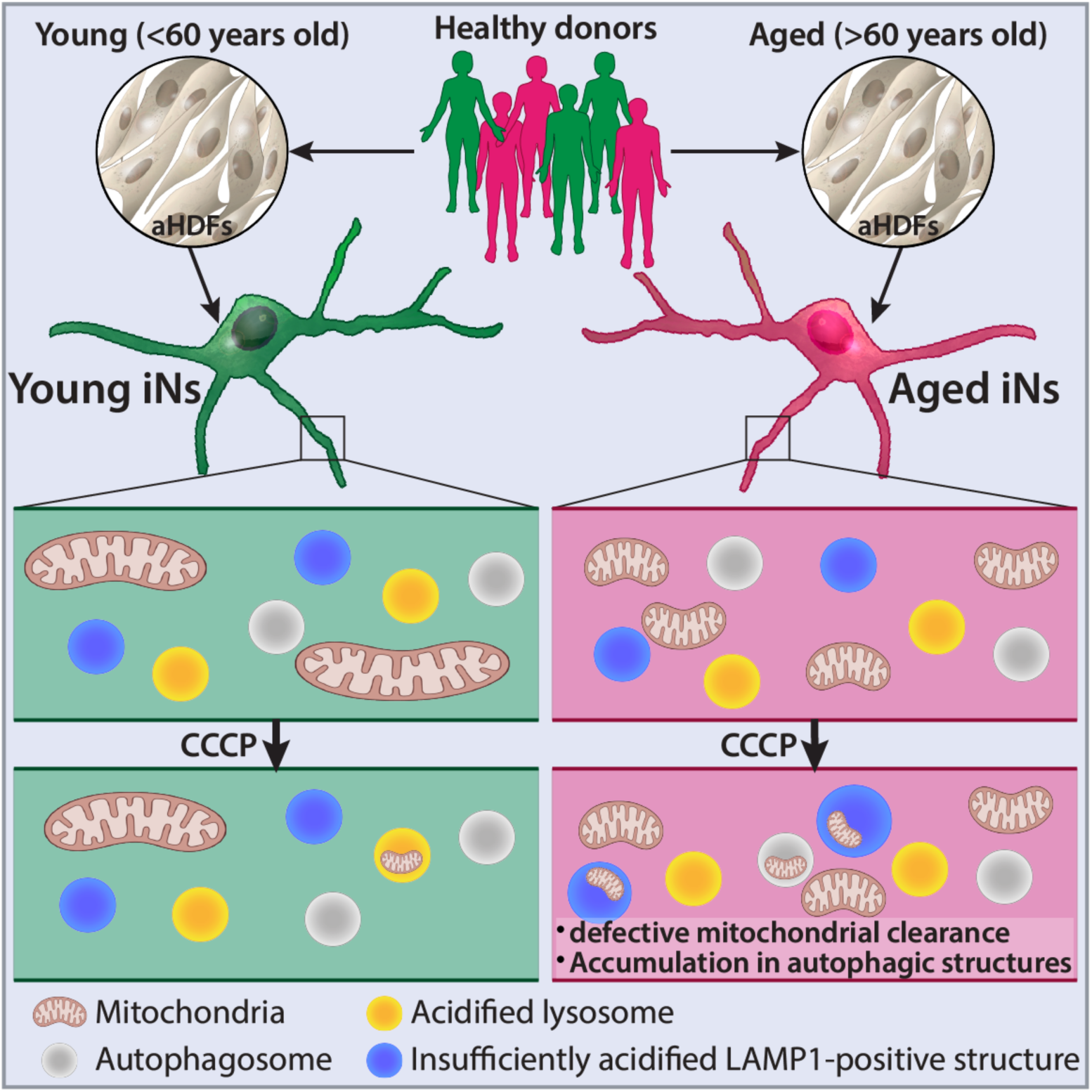

## 1. Introduction

Aging is a major risk factor for many neurodegenerative diseases, including PD and AD (Y. Hou et al., 2019). However, how age-related molecular processes lead to neurodegeneration in human neurons is still unclear(Rhine et al., 2025). To date, most of our knowledge on how normal aging affects the brain, and more specifically neurons, comes from animal models. Studies in rodents have shown that normal brain aging at a cellular level is associated with, amongst other, neuronal protein aggregation(Cuanalo-Contreras et al., 2023; Guldner et al., 2026), impairment of the main protein degradation systems (ubiquitin-proteasome system (UPS)(Keller et al., 2000; Parker et al., 2025) and autophagy(Yang et al., 2014; Zheng et al., 2021)), nuclear pore integrity(Rempel et al., 2020; Savas et al., 2012), lysosomal dysfunction(Burrinha et al., 2023; Nakanishi et al., 1994; Nixon, 2020), as well as increased oxidative damage(Aksenova et al., 1998; Çakatay et al., 2001; Floyd & Hensley, 2002; Hamilton et al., 2001). These deficits have also been linked to neurodegeneration(Burrinha et al., 2023; Kluever & Fornasiero, 2021; Lee & Kim, 2022; Palmer et al., 2025). Other common non-murine aging models such as *Caenorhabditis elegans* (*C. elegans*) and *Drosophila melanogaster* (*D. melanogaster*), and less common ones such as non-human primates, partially recapitulate those findings(Loeffler, 2019).

While animal models are greatly contributing to our current understanding of how aging affects fundamental cellular processes in neurons, they present some limitations –an important one being their considerably shorter lifespan compared to the human lifespan (Kluever & Fornasiero, 2021). This difference is particularly relevant for post-mitotic cells, such as neurons. Neurons, in most regions of the central nervous system stop dividing early during development, and their population is not substantially replenished throughout the lifespan. As a result, damaged cell components and protein aggregates can progressively accumulate over the course of the life of the cell and can lead to increased cellular damage(Carmona-Gutierrez et al., 2016; Rubinsztein et al., 2011; Terman et al., 2010). Their shorter lifespan might contribute to the reason why animals generally do not spontaneously develop age-associated neurodegenerative diseases(Jucker, 2010), therefore only allowing for the modelling of the less common genetic forms of diseases such as AD and PD (Link, 2001). Furthermore, lower inter-individual heterogeneity due to inbreeding (Cohen, 2018; Flurkey & Currer, 2004) and inter-species intrinsic biological differences in molecular mechanisms of aging (Cohen, 2018) are other limitations associated with the use of animal models in the context of aging studies. Although most of the current knowledge on cellular mechanisms of neuronal aging –both in normal aging and pathological conditions –relies on animal studies, a limited number of human post-mortem brain studies of normal aging are available. Similarly to what has been reported in animal models, these studies show, with advancing age, decreased autophagy(Lipinski et al., 2010; Loeffler, 2019; Shibata et al., 2006), decreased mitochondrial health and function(Bender et al., 2006; Cortopassi & Arnheim, 1990; Cottrell et al., 2001; X. Hou et al., 2018; Itoh et al., 1996; Kennedy et al., 2013; Kraytsberg et al., 2006; Venkateshappa et al., 2012), increased overall protein oxidation(Smith et al., 1991), oxidative stress(Tokarew et al., 2021; Venkateshappa et al., 2012), and neurodegeneration(Fearnley & Lees, 1991). While these findings in human brain tissue are important to pin-point age-related changes in human neurons, they only provide a snapshot, rather than a dynamic examination of molecular process impairment over time. Induced neurons (iNs) generated through direct neuronal reprogramming of dermal fibroblasts from human donors are currently one of the few elective methods to overcome these limitations, as they offer a platform to study the effect of aging dynamically and over time(Bury et al., 2025). One key advantage of this tool is the maintenance of age-related signatures(Mertens et al., 2018) related to mitochondrial health and function(Kim et al., 2018; Varghese et al., 2025), nucleocytoplasmic compartmentalization(Mertens et al., 2015), autophagy (Drouin-Ouellet et al., 2022), oxidative damages and oxidative stress levels(Huh et al., 2016; Kim et al., 2018), DNA damage(Chou et al., 2025; Drouin-Ouellet et al., 2022; Huh et al., 2016), DNA methylation marks(Chou et al., 2025; Huh et al., 2016; Mertens et al., 2021; Pircs et al., 2021), cellular stress and localization of splicing proteins(Rhine et al., 2025). Interestingly, the findings in iNs point towards age-related impairment in the same cellular pathways as that reported in human post-mortem brain samples, supporting the usefulness of this model to study age-related human neuronal changes.

Mitochondrial dysfunction is often considered a hallmark of aging(López-Otín et al., 2013; Sun et al., 2016). Mitochondria are crucial organelles for metabolic support, especially in highly metabolic cells that are mostly relying on oxidative phosphorylation such as neurons(Area-Gomez et al., 2019; Kim et al., 2018; Mahadevan et al., 2021). As such, their proper functioning is central for neuronal survival, neurotransmitter release, axonal transport and neuroplasticity(Kann & Kovács, 2007; Mahadevan et al., 2021). As previously mentioned, mitochondrial health decreases in healthy aging, as observed in human post-mortem brain samples (Bender et al., 2006; Cortopassi & Arnheim, 1990; Cottrell et al., 2001; X. Hou et al., 2018; Itoh et al., 1996; Kennedy et al., 2013; Kraytsberg et al., 2006; Venkateshappa et al., 2012) and iNs (Kim et al., 2018; Varghese et al., 2025). Identified impairments include decreased mitochondrial membrane potential (ΔΨm)(Kim et al., 2018; Varghese et al., 2025), decreased electron transport chain (ETC) complex IV activity(Cottrell et al., 2001; Itoh et al., 1996), increased mitochondrial network fragmentation(Kim et al., 2018; Varghese et al., 2025), decreased energy production(Kim et al., 2018; Varghese et al., 2025), increased reactive oxygen species (ROS) production(Kim et al., 2018; Smith et al., 1991; Varghese et al., 2025; Venkateshappa et al., 2012), accumulation of mitochondrial DNA mutations(Bender et al., 2006; Cortopassi & Arnheim, 1990; Kennedy et al., 2013; Kraytsberg et al., 2006) and decreased mitochondrial genes expression(Kim et al., 2018). Such mitochondrial alterations lead to aberrant generation of ROS(Banarase et al., 2023). In turn, excessive ROS induce oxidative damage to lipids, DNA and proteins, increasing mitochondrial DNA damage and ultimately leading to cellular dysfunction and death(Banarase et al., 2023; Golpich et al., 2017). The efficient elimination of damaged mitochondria via mitophagy could contribute to breaking this cycle and preserve neuronal health during aging(Mishra & Thakur, 2023; Schmid et al., 2022).

Damaged mitochondria are mainly eliminated by autophagy –a process called mitophagy –mostly by ubiquitin-dependant pathways such as the PINK1/Parkin-dependent pathway, and in part by ubiquitin-independent pathways involving BNIP3, BNIP3L/NIX and FUNDC1(Ganley & Simonsen, 2022; S. Wang et al., 2023). Mitophagy defects have been associated with age-related neurodegenerative diseases such as PD and AD(Cai & Jeong, 2020) and with normal aging in numerous animal models(Rappe & McWilliams, 2022). However, despite the widely accepted role of mitochondrial decline in aging, and evidence of age-associated autophagy defects, the molecular mechanisms underlying impaired mitophagy in human neuronal aging remain poorly characterized. Understanding how mitophagy progressively decreases during aging is crucial to devise strategies to promote healthy aging and to determine the implications of this process in age-related neurodegenerative diseases.

Here, by comparing iNs reprogrammed from dermal fibroblasts derived from young (<60 years old) and aged (>60 years old) healthy donors, we replicate earlier findings in iNs showing age-associated decrease in ΔΨm and increased mitochondrial fragmentation. We further show age-related mitophagy impairment, which are characterized by an accumulation of mitochondria in autophagic structures leading to a decline in mitochondrial clearance. Importantly, we identify defective translocation of mitochondria to acidified lysosomes as the likely step preventing effective mitochondrial clearance in aging. By reporting for the first time age-related mitophagy impairments in human neurons, this study opens the path for future mechanistic studies and the identification of novel strategies to boost mitochondrial clearance in aging neurons.

## 2. Material and methods

### 2.1. Human fibroblasts

Adult human dermal fibroblasts (aHDFs) derived from young (<60 years old) and aged (>60 years old) donors without any reported medical condition were acquired from the Coriell Institute (see **Table 1**). Fibroblasts were cultured in T75 culture flasks in DMEM (high glucose, with sodium pyruvate and GlutaMAX) supplemented with 10% fetal bovine serum and 100U/mL of penicillin/streptomycin (P/S) at 37°C in 5% CO_2_. Cells were passaged when reaching ∼80% confluence. For passaging and plating for neuronal conversion, cells were incubated with 0.05% trypsin-EDTA for 5 min at 37°C in 5% CO_2_ before being resuspended and collected in fresh medium. aHDFs were spun down at 400g for 5 min, supernatant was then discarded, and aHDFs were resuspended in fresh medium. A Countess™ automated cell counter was used to determine viable cell number for plating. All cell lines were tested for mycoplasma before plating for neuronal conversion using a PCR mycoplasma detection kit, and were negative for mycoplasma. For details on materials and catalog numbers, refer to **Table 2**.

**Table 1.** Cell lines.

| Category | Identifier | Age | Sex | Race | Skin Biopsy Location | Source | RRID |
| --- | --- | --- | --- | --- | --- | --- | --- |
| Young (<60 years old) | GM02037 | 13 | Male | White | Unspecified | Coriell Institute | CVCL_7349 |
|  | AG11747 | 22 | Male | White | Mesial aspect of the mid-upper left arm | Coriell Institute | CVCL_2E46 |
|  | AG09860 | 26 | Male | White | Mesial aspect of the mid-upper left arm | Coriell Institute | CVCL_2D20 |
|  | AG13011 | 35 | Male | White | Mesial aspect of the mid-upper left arm | Coriell Institute | CVCL_2F26 |
|  | AG05805 | 52 | Male | White | Mesial aspect of the mid-upper left arm | Coriell Institute | CVCL_2B53 |
|  | GM01652 | 11 | Female | White | Left forearm | Coriell Institute | CVCL_7330 |
|  | AG07720 | 24 | Female | White | Mesial aspect of the mid-upper left arm | Coriell Institute | CVCL_2C42 |
|  | AG07306 | 28 | Female | White | Mesial aspect of the mid-upper left arm | Coriell Institute | CVCL_2C28 |
|  | AG07121 | 34 | Female | White | Mesial aspect of the mid-upper left arm | Coriell Institute | CVCL_2C16 |
|  | AG09162 | 55 | Female | White | Mesial aspect of the mid-upper left arm | Coriell Institute | CVCL_2C92 |
| Aged (>60 years old) | AG07996 | 65 | Male | White | Mesial aspect of the mid-upper left arm | Coriell Institute | CVCL_2C51 |
|  | AG13108 | 75 | Male | White | Mesial aspect of the mid-upper left arm | Coriell Institute | CVCL_2F54 |
|  | AG06243 | 82 | Male | White | Mesial aspect of the mid-upper left arm | Coriell Institute | CVCL_2B73 |
|  | AG07725 | 91 | Male | White | Mesial aspect of the mid-upper left arm | Coriell Institute | CVCL_2C46 |
|  | AG11357 | 62 | Female | White | Mesial aspect of the mid-upper left arm | Coriell Institute | CVCL_2D90 |
|  | AG09225 | 72 | Female | White | Mesial aspect of the mid-upper left arm | Coriell Institute | CVCL_2C94 |
|  | AG13077 | 85 | Female | White | Mesial aspect of the mid-upper left arm | Coriell Institute | CVCL_2F33 |
|  | AG08712 | 90 | Female | White | Forearm | Coriell Institute | CVCL_4P83 |

**Table 2.**
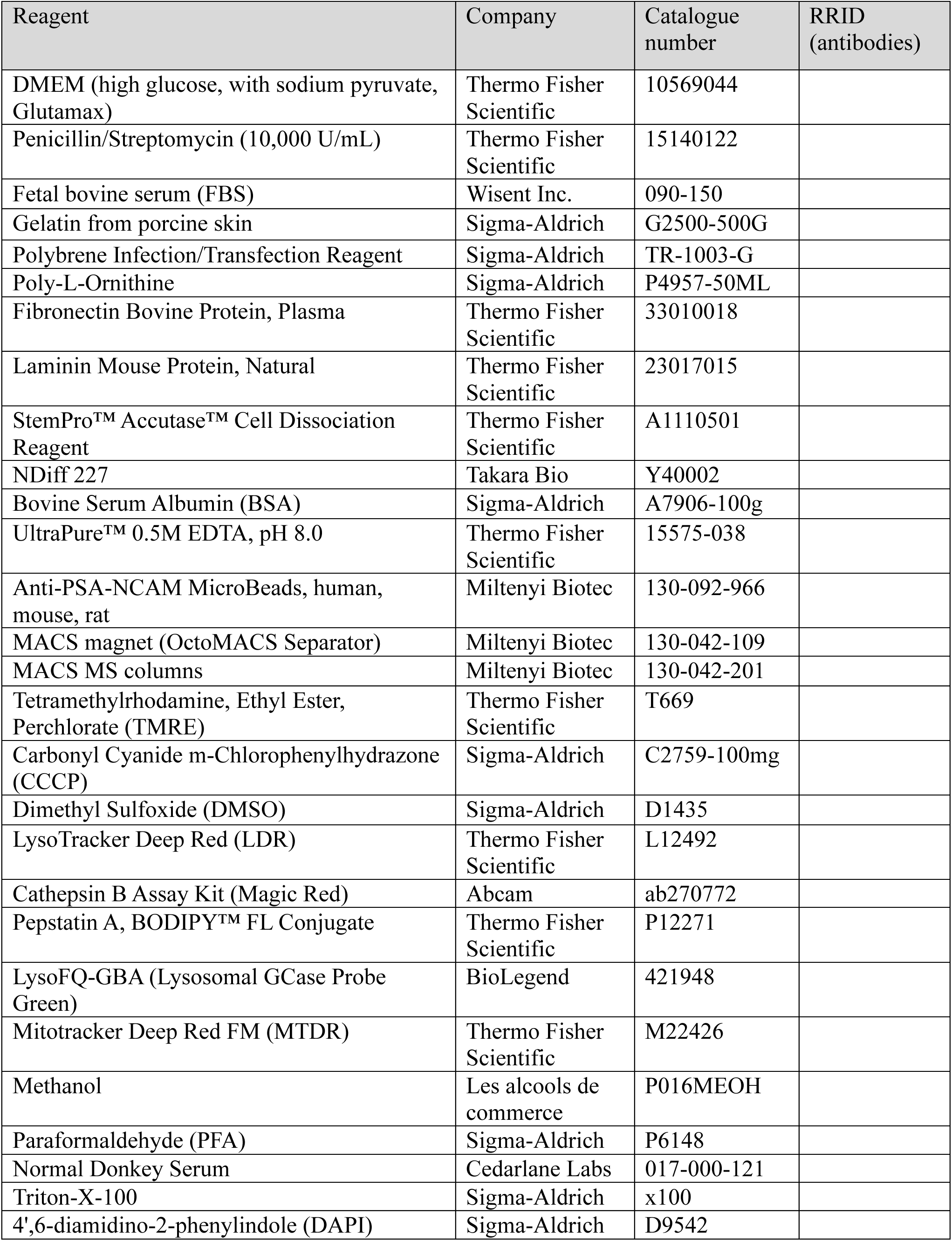
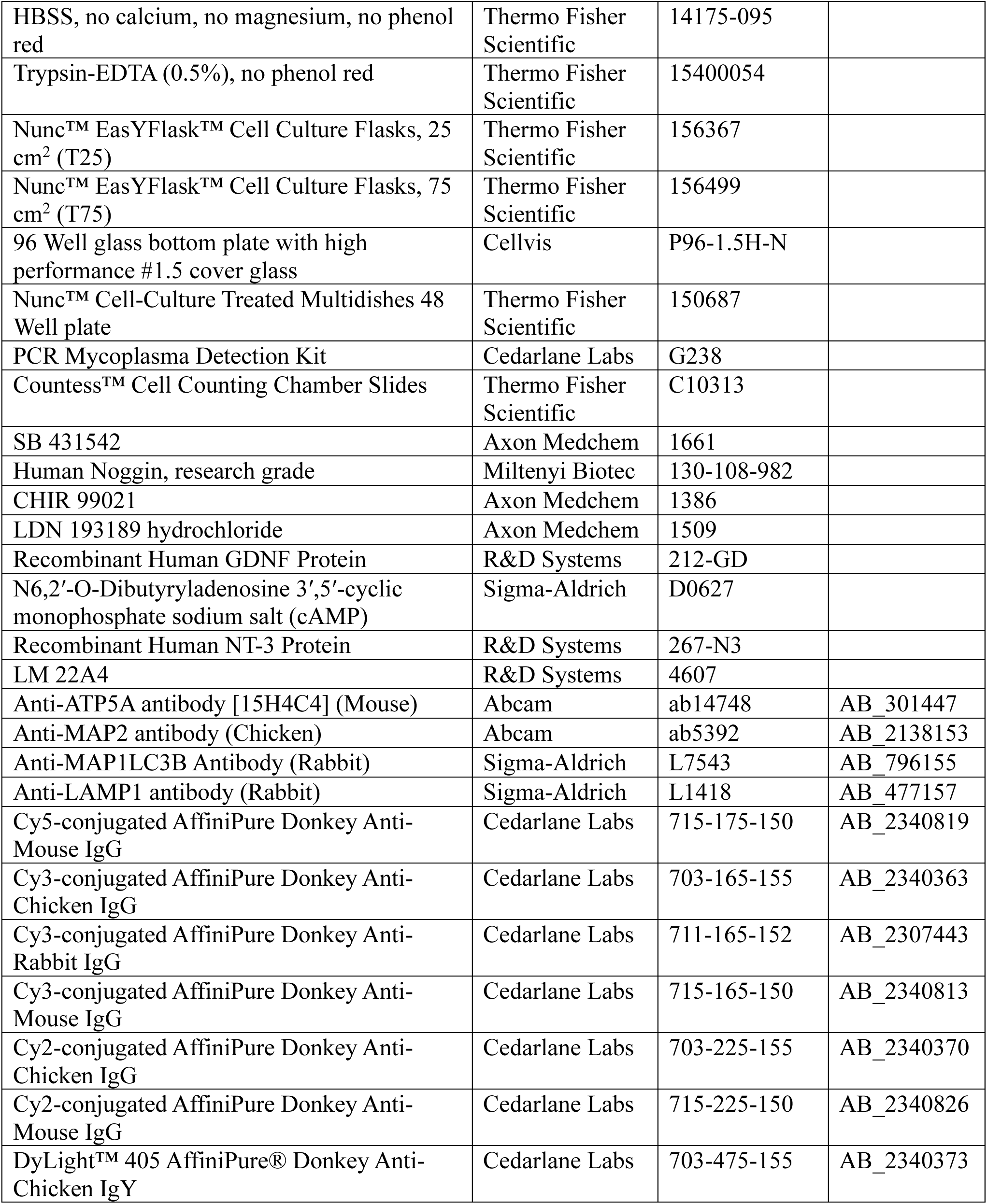
List of materials, reagents and antibodies.

| Reagent | Company | Catalogue number | RRID (antibodies) |
| --- | --- | --- | --- |
| DMEM (high glucose, with sodium pyruvate, Glutamax) | Thermo Fisher Scientific | 10569044 |  |
| Penicillin/Streptomycin (10,000 U/mL) | Thermo Fisher Scientific | 15140122 |  |
| Fetal bovine serum (FBS) | Wisent Inc. | 090-150 |  |
| Gelatin from porcine skin | Sigma-Aldrich | G2500-500G |  |
| Polybrene Infection/Transfection Reagent | Sigma-Aldrich | TR-1003-G |  |
| Poly-L-Ornithine | Sigma-Aldrich | P4957-50ML |  |
| Fibronectin Bovine Protein, Plasma | Thermo Fisher Scientific | 33010018 |  |
| Laminin Mouse Protein, Natural | Thermo Fisher Scientific | 23017015 |  |
| StemPro™ Accutase™ Cell Dissociation Reagent | Thermo Fisher Scientific | A1110501 |  |
| NDiff 227 | Takara Bio | Y40002 |  |
| Bovine Serum Albumin (BSA) | Sigma-Aldrich | A7906-100g |  |
| UltraPure™ 0.5M EDTA, pH 8.0 | Thermo Fisher Scientific | 15575-038 |  |
| Anti-PSA-NCAM MicroBeads, human, mouse, rat | Miltenyi Biotec | 130-092-966 |  |
| MACS magnet (OctoMACS Separator) | Miltenyi Biotec | 130-042-109 |  |
| MACS MS columns | Miltenyi Biotec | 130-042-201 |  |
| Tetramethylrhodamine, Ethyl Ester, Perchlorate (TMRE) | Thermo Fisher Scientific | T669 |  |
| Carbonyl Cyanide m-Chlorophenylhydrazone (CCCP) | Sigma-Aldrich | C2759-100mg |  |
| Dimethyl Sulfoxide (DMSO) | Sigma-Aldrich | D1435 |  |
| LysoTracker Deep Red (LDR) | Thermo Fisher Scientific | L12492 |  |
| Cathepsin B Assay Kit (Magic Red) | Abcam | ab270772 |  |
| Pepstatin A, BODIPY™ FL Conjugate | Thermo Fisher Scientific | P12271 |  |
| LysoFQ-GBA (Lysosomal GCase Probe Green) | BioLegend | 421948 |  |
| Mitotracker Deep Red FM (MTDR) | Thermo Fisher Scientific | M22426 |  |
| Methanol | Les alcools de commerce | P016MEOH |  |
| Paraformaldehyde (PFA) | Sigma-Aldrich | P6148 |  |
| Normal Donkey Serum | Cedarlane Labs | 017-000-121 |  |
| Triton-X-100 | Sigma-Aldrich | x100 |  |
| 4',6-diamidino-2-phenylindole (DAPI) | Sigma-Aldrich | D9542 |  |
| HBSS, no calcium, no magnesium, no phenol red | Thermo Fisher Scientific | 14175-095 |  |
| Trypsin-EDTA (0.5%), no phenol red | Thermo Fisher Scientific | 15400054 |  |
| Nunc™ EasYFlask™ Cell Culture Flasks, 25 cm<sup>2</sup> (T25) | Thermo Fisher Scientific | 156367 |  |
| Nunc™ EasYFlask™ Cell Culture Flasks, 75 cm<sup>2</sup> (T75) | Thermo Fisher Scientific | 156499 |  |
| 96 Well glass bottom plate with high performance #1.5 cover glass | Cellvis | P96-1.5H-N |  |
| Nunc™ Cell-Culture Treated Multidishes 48 Well plate | Thermo Fisher Scientific | 150687 |  |
| PCR Mycoplasma Detection Kit | Cedarlane Labs | G238 |  |
| Countess™ Cell Counting Chamber Slides | Thermo Fisher Scientific | C10313 |  |
| SB 431542 | Axon Medchem | 1661 |  |
| Human Noggin, research grade | Miltenyi Biotec | 130-108-982 |  |
| CHIR 99021 | Axon Medchem | 1386 |  |
| LDN 193189 hydrochloride | Axon Medchem | 1509 |  |
| Recombinant Human GDNF Protein | R&D Systems | 212-GD |  |
| N6,2'-O-Dibutyryl adenosine 3',5'-cyclic monophosphate sodium salt (cAMP) | Sigma-Aldrich | D0627 |  |
| Recombinant Human NT-3 Protein | R&D Systems | 267-N3 |  |
| LM 22A4 | R&D Systems | 4607 |  |
| Anti-ATP5A antibody [15H4C4] (Mouse) | Abcam | ab14748 | AB_301447 |
| Anti-MAP2 antibody (Chicken) | Abcam | ab5392 | AB_2138153 |
| Anti-MAP1LC3B Antibody (Rabbit) | Sigma-Aldrich | L7543 | AB_796155 |
| Anti-LAMP1 antibody (Rabbit) | Sigma-Aldrich | L1418 | AB_477157 |
| Cy5-conjugated AffiniPure Donkey Anti-Mouse IgG | Cedarlane Labs | 715-175-150 | AB_2340819 |
| Cy3-conjugated AffiniPure Donkey Anti-Chicken IgG | Cedarlane Labs | 703-165-155 | AB_2340363 |
| Cy3-conjugated AffiniPure Donkey Anti-Rabbit IgG | Cedarlane Labs | 711-165-152 | AB_2307443 |
| Cy3-conjugated AffiniPure Donkey Anti-Mouse IgG | Cedarlane Labs | 715-165-150 | AB_2340813 |
| Cy2-conjugated AffiniPure Donkey Anti-Chicken IgG | Cedarlane Labs | 703-225-155 | AB_2340370 |
| Cy2-conjugated AffiniPure Donkey Anti-Mouse IgG | Cedarlane Labs | 715-225-150 | AB_2340826 |
| DyLight™ 405 AffiniPure® Donkey Anti-Chicken IgY | Cedarlane Labs | 703-475-155 | AB_2340373 |

### 2.2. Direct neuronal reprogramming from human fibroblasts

For most experiments, aHDFs were plated in either T25 (300K cells) or T75 flasks (1-2 million cells), previously coated overnight with 0.1% gelatin. For experiments related to **Fig. 4**, aHDFs were plated in 96 well plates (7,500 cells per well), previously coated with a combination of polyornithine (15 μg/mL), fibronectin (0.5 ng/μL), and laminin (5 μg/mL)(Shrigley et al., 2018). A third-generation lentivirus vector containing ASCL1, BRN2 and shRNA sequences against REST(Drouin-Ouellet et al., 2017; Shrigley et al., 2018) under a non-regulated ubiquitous phosphoglycerate kinase (PGK) promoter was produced as previously described(Drouin-Ouellet et al., 2017; Shrigley et al., 2018). Fibroblasts were directly reprogrammed into iNs using a previously published protocol(Shrigley et al., 2018). Briefly, transduction was done in medium containing 8µg/mL polybrene. Cells were differentiated in early neuronal medium (ENM), and the medium was replaced by late neuronal medium (LNM) for neuronal maturation at day 18 of conversion(Drouin-Ouellet et al., 2017). See **Table 2** and (Shrigley et al., 2018) for media composition, and **Fig. 1a** for a schematic representation of the experimental workflow. A total of 4 independent reprogramming experiments were conducted. For the reprogramming experiments leading to TMRE and LC3 results, one cell line (75, M) failed to reprogram, and therefore this cell line was excluded from the analysis in **Fig. 1b-e** and **Fig. 2a-c**. For the reprogramming experiments leading to the LAMP1 results, one cell line (28, F) also failed to reprogram, and therefore is excluded from **Fig. 2d-h**. For the lysosomal enzyme activity experiments (**Fig. 4**), one cell line (72, F) was not included in the conversion due to exhaustion of the available cell stock.

**Figure 1.**
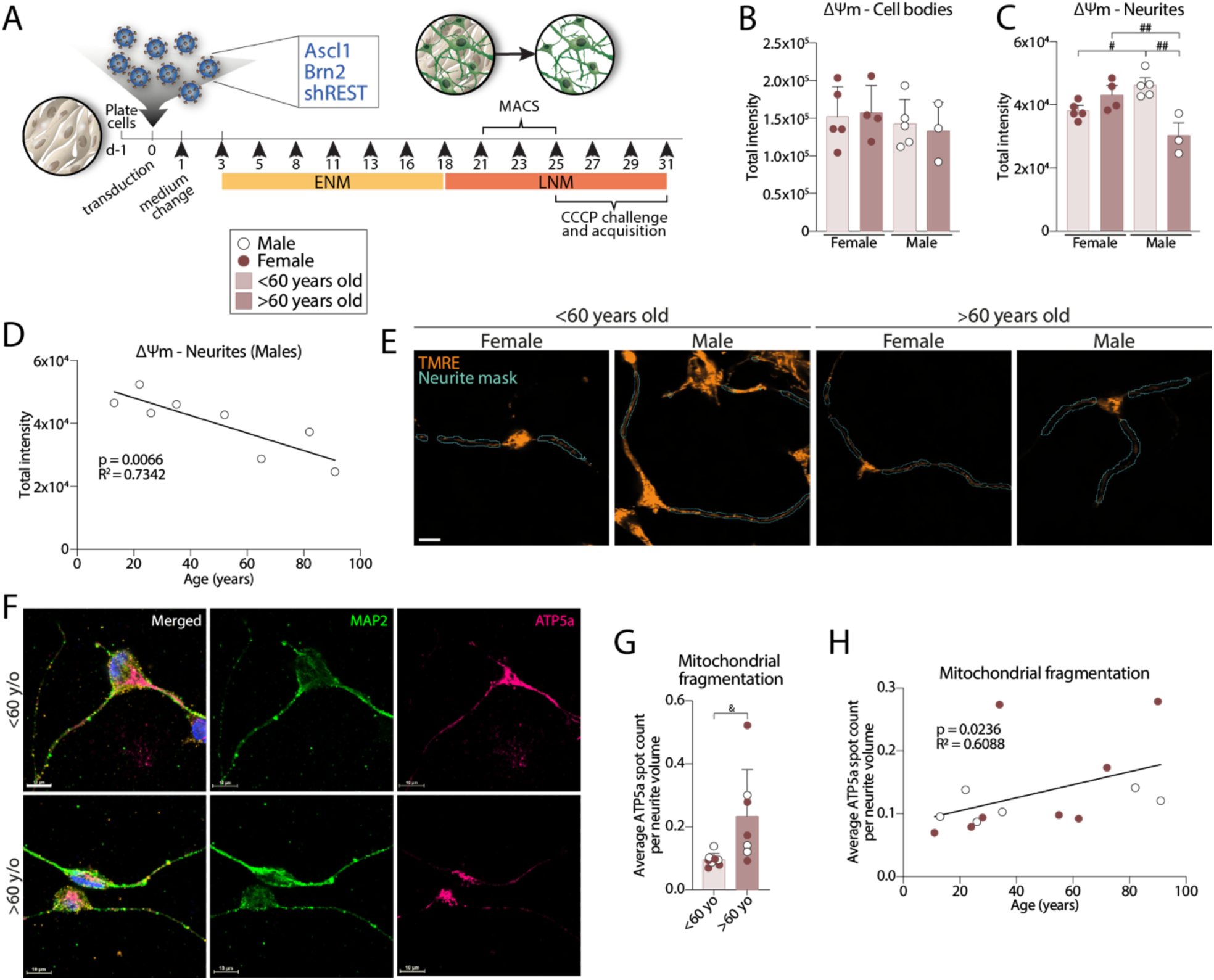
Decreased basal mitochondrial health in neurites with healthy aging. **(A)** Schematic representation of the main steps of the reprogramming protocol. **(B-C)** Total intensity of TMRE in cell bodies (A) and neurites (B) of iNs. Neurites: One-way ANOVA, Tukey’s multiple comparisons test; males, ##*p*=0.0011; under 60 years old, #*p*=0.0484; over 60 years old, ##*p*=0.0088. Data are expressed as mean ± the SD. **(D)** Correlation between age and average total intensity of TMRE per object in neurites of iNs derived from male donors. Pearson’s correlation rank: p=0.0066; 95% confidence interval -0.9736 to -0.3841. **(E)** Representative fluorescence images of TMRE staining, and representative neurite segmentation in iNs derived from male and female donors aged under or over 60 years old. **(F)** Representative immunofluorescence images of the neuronal marker MAP2 and the mitochondrial marker ATP5a in iNs of donors aged under and over 60 years old. **(G)** Quantification of the average ATP5a puncta count per neurite volume in iNs derived from both age groups. Two-tailed unpaired t-test: &p=0.0230, df=13. Data are expressed as mean ± the SD. **(H)** Correlation between the average ATP5a puncta count per neurite volume and age of the donors (n= 14 donors). Spearman’s rank correlation: p=0.0236; 95% confidence interval 0.09826 to 0.8656. Abbreviations: ΔΨm, mitochondrial membrane potential; TMRE, Tetramethylrhodamine ethyl ester; yo, years old.

**Figure 2.**
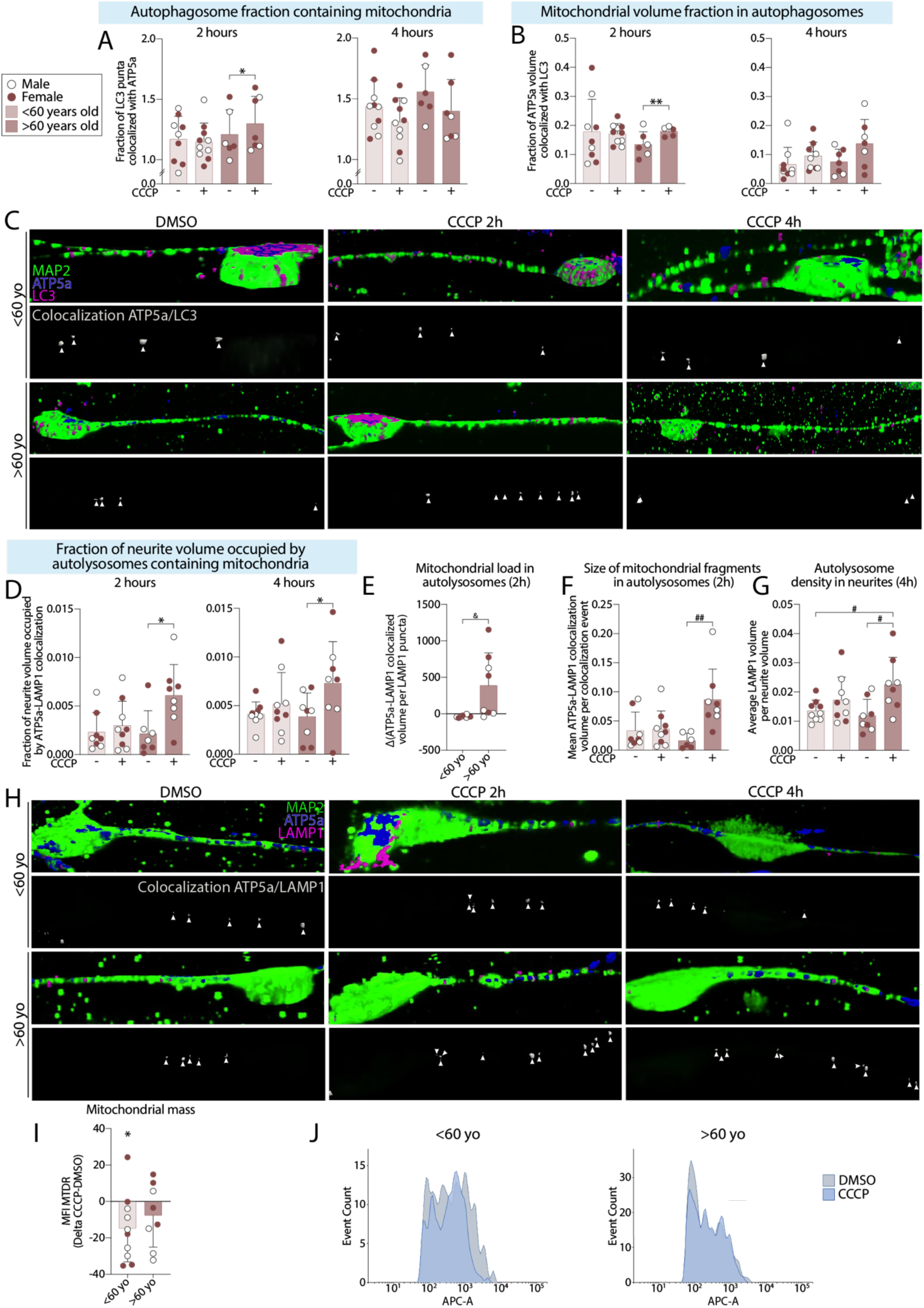
Age-associated mitophagy impairment in neurites. **(A)** Quantification of the fraction (by count) of LC3 puncta positive for ATP5a after 2 and 4 hours of 20µM CCCP challenge. 2 hours: Two-tailed paired t-test, \**p*=0.0205, df=5. Data are expressed as mean ± the SD. **(B)** Quantification of the fraction (by volume) of ATP5a puncta positive for LC3 after 2 and 4 hours of CCCP challenge. 2 hours: Two-tailed paired t-test, \*\**p*=0.0036, df=3. Data are expressed as mean ± the SD. **(C)** Representative 3D reconstruction of the immunofluorescence images of MAP2, ATP5a and LC3 after exposure to CCCP for 2 and 4 hours, as well as representative 3D reconstruction of the ATP5a-LC3 puncta colocalization in the MAP2-positive neurites. **(D)** Quantification of the fraction of neurite volume occupied by of ATP5a and LAMP1 double-positive puncta at 2 and 4 hours of CCCP challenge. 2 hours: over 60 years old, Wilcoxon matched pairs signed rank test, \**p*=0.0312, rs=0.3095. 4 hours: over 60 years old, two-tailed paired t-test, \**p*=0.0131, df=7. Data are expressed as mean ± the SD. **(E)** Quantification of the fold change of the ATP5a-LAMP1 colocalized volume per LAMP1 puncta in neurites CCCP exposure for 2 hours. Two-tailed unpaired t-test, &*p*=0.0261, df=13. Data are expressed as mean of the fold change induced by CCCP compared to the vehicle ± the SD. **(F)** Quantification of the mean ATP5a-LAMP1 double-positive puncta volume per colocalization event following 2 hours of CCCP exposure. Over 60 years old, Kruskal-Wallis test, Dunn’s multiple comparisons test, ##*p*=0.0058. Data are expressed as mean ± the SD. **(G)** Quantification of the average LAMP1 volume per neurite volume after a 4 hour CCCP exposure. One-way ANOVA, Tukey’s multiple comparisons test: over 60 years old, #*p*=0.0189; under 60 years old vehicle and over 60 years old CCCP, #*p*=0.0498. Data are expressed as mean ± the SD. **(H)** Representative 3D reconstruction of the immunofluorescence images of MAP2, ATP5a and LAMP1 after exposure to vehicle, or CCCP for 2 hours and 4 hours, as well as representative 3D reconstruction of the ATP5a-LAMP1 puncta colocalization in the MAP2-positive neurites. **(I)** Quantification of the fold change in the mean fluorescence intensity of Mitotracker Deep Red FM induced by exposure to CCCP for 4 hours. Under 60 years old: two-tailed paired t-test, \**p*=0.0369, df=9. Data are expressed as mean of the fold change induced by CCCP compared to the vehicle ± the SD. **(J)** Representative histogram of flow cytometry analysis of Mitotracker Deep Red FM staining in iNs after exposure to vehicle or CCCP for 4 hours. Data are expressed as mean ± the SD. Abbreviations: CCCP, Carbonyl cyanide 3-chlorophenylhydrazone; MFI, mean fluorescence intensity; MTDR, Mitotracker Deep Red FM; yo, years old.

### 2.3. iN purification and replating

For experiments related to **Fig. 1-3**, between day 21 and day 25 of conversion, iNs were purified by magnetic-activated cell sorting (MACS).

**Figure 3.**
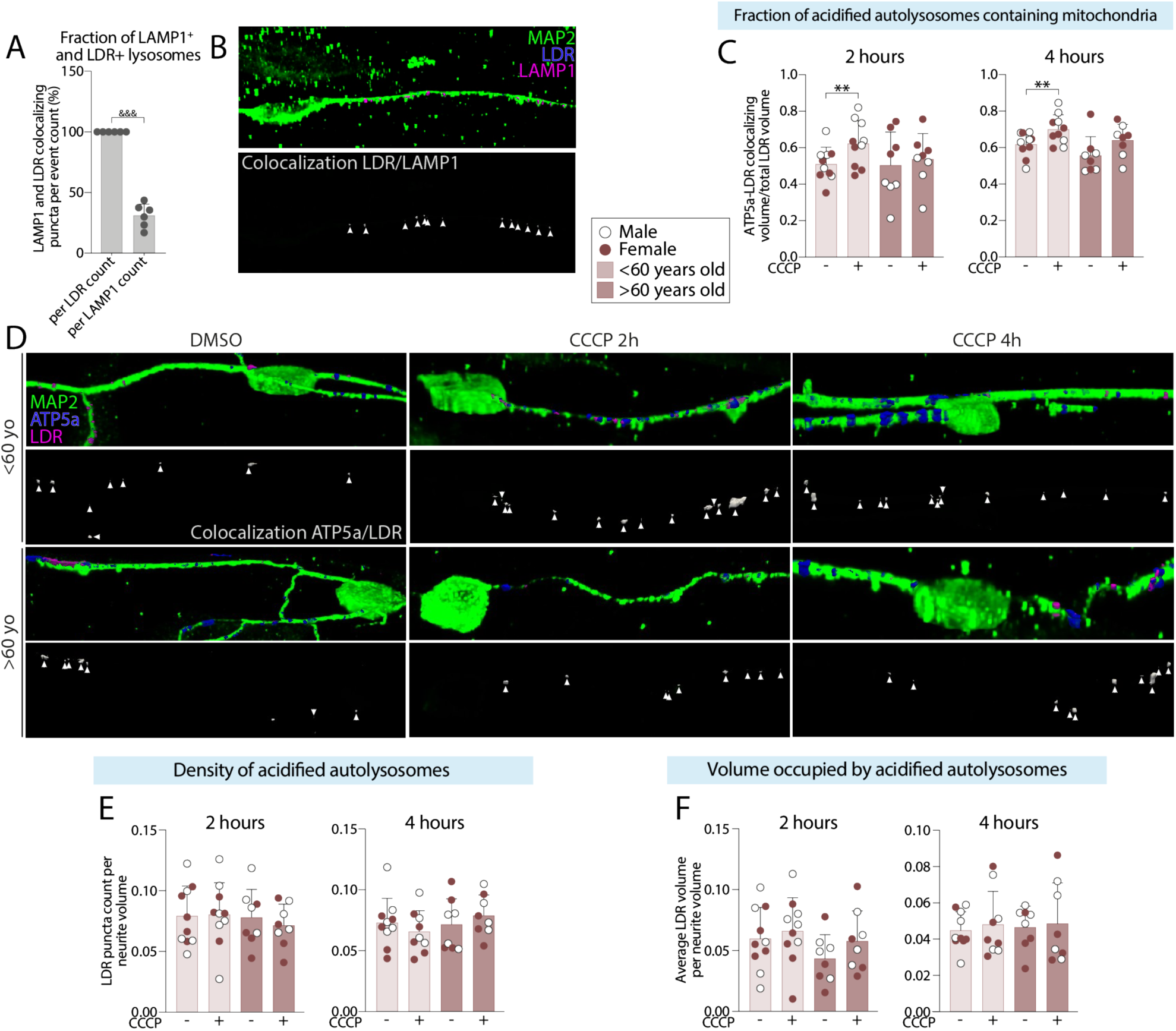
Age-associated impairment in translocation of mitochondria to acidified autolysosomes in neurites. **(A)** Quantification of the fraction (by count) of LAMP1 and Lysotracker Deep Red double-positive puncta per Lysotracker Deep Red puncta count or by LAMP1 puncta count. Two-tailed unpaired t-test, &&&*p*<0.0001, df=10. Data are expressed as mean ± the SD. **(B)** Representative 3D reconstruction of the immunofluorescence images of MAP2, Lysotracker Deep Red and LAMP1 staining, as well as representative 3D reconstruction of the Lysotracker Deep Red and LAMP1 double-positive puncta in the MAP2-positive neurites. **(C)** Quantification of the fraction (by volume) of Lysotracker Deep Red puncta positive for ATP5a after 2 and 4 hours of CCCP exposure. 2 hours: two-tailed paired t-test, \*\**p*=0.0029, df=9. 4 hours: under 60 years old, two-tailed paired t-test, \*\**p*=0.0018, df=9; vehicles. Data are expressed as mean ± the SD. **(D)** Representative 3D reconstruction of the immunofluorescence images of MAP2, ATP5a and Lysotracker Deep Red, as well as representative 3D reconstruction of the ATP5a and Lysotracker Deep Red double-positive puncta in the MAP2-positive neurites. **(E)** Quantification of the density (by count) of Lysotracker Deep Red-positive autolysosomes per neurite volume after 2 and 4 hours of CCCP exposure. Data are expressed as mean ± the SD. **(F)** Quantification of the density (by volume) of Lysotracker Deep Red-positive autolysosome per neurite volume after 2 and 4 hours of CCCP exposure. Data are expressed as mean ± the SD. Abbreviations: CCCP, Carbonyl cyanide 3-chlorophenylhydrazone; DMSO, Dimethyl sulfoxide; LDR, Lysotracker Deep Red; yo, years old.

Cells were incubated for 5-10 min with Accutase at 37°C in 5% CO_2_, then collected in NDiff227 medium containing 1% P/S. After centrifugation of the cell suspension for 5 min at 400g, the pellet was resuspended in a mix of 20µL LNM+20µL DPBS 1X containing 0.5% BSA and 2mM EDTA, with 10µL anti-PSA-NCAM microbeads, and incubated for 15 min at 4°C. During the incubation, the MACS columns (while on the magnet) were equilibrated with 500µL NDiff227 + 1% P/S. After the incubation, 500µL of NDiff227 + 1% P/S was added to the cells before centrifugation (400g, 5 min). The pellet was resuspended in 500µL NDiff227 + 1% P/S and run through the column. The column was washed 3 times with 500µL NDiff227 + 1% P/S before being removed from the magnet. The iNs were then eluted in 500µL LNM. The live iNs were counted using a Countess™ automatic cell counter, spun down (400g, 5 min) and resuspended at a concentration of 250 cells/µL in LNM for seeding. Cells were seeded as a sitting drop for 1h at 37°C in 5% CO_2_ on multi-well plates previously coated with a combination of polyornithine (15 μg/mL), fibronectin (0.5 ng/μL), and laminin (5 μg/mL)(Shrigley et al., 2018), before additional medium was added to the well. For microscopy analysis, 5000 iNs per well were replated in a sitting drop in 96 wells glass-bottom plates. For flow cytometry analysis, 15,000 iNs per well were replated in 48 well plates. iNs were maintained in LNM, with half medium changes every 2-3 days until the end of experiment. Challenges and analysis were conducted between day 25 and 31 of conversion, minimally 4 days after MACS, with the exception of LysoFQ-GBA analysis, which was conducted at day 35 of conversion. The conversion efficiency of iNs can be found in **Supp.Fig. 1a**, as well as a representative image of the resulting purified culture (**Supp. Fig. 1b**).

### 2.4. Measurement of mitochondrial membrane potential

Mitochondrial membrane potential (ΔΨm) was assessed using Tetramethylrhodamine ethyl ester (TMRE) probe. TMRE was resuspended in dimethyl sulfoxide (DMSO) to obtain a stock concentration of 10mM, and stored at -20°C. To assess ΔΨm, iNs were incubated for 30 min at 37°C in 5% CO_2_ in LNM containing 5nM TMRE. Cells were then washed once with DPBS 1X, and live images were acquired in LNM in a controlled environment chamber (Tokai Hit stage top incubation system, 37°C, 5% CO_2_).

### 2.5. Mitophagy induction

Mitophagy was induced by mitochondrial depolarization using carbonyl cyanide m-chlorophenyl hydrazone (CCCP). CCCP was resuspended in DMSO to obtain a stock concentration of 100mM, and stored at -20°C. iNs were exposed to a solution of 20 µM of CCCP or a corresponding volume of DMSO in LNM for 2 or 4h (37°C in 5% CO_2_)(Iriondo et al., 2022; Tanaka et al., 2010).

### 2.6. Lysotracker Deep Red staining

Lysotracker Deep Red (LDR) was diluted to a working solution of 100nM in LNM. Following the indicated treatments, iNs were incubated with the working solution for 30 min at 37°C in 5% CO_2_. Medium was then removed, cells were washed twice with DPBS 1X, then fixed with 4% ice-cold paraformaldehyde (PFA) for 10 min. PFA was then removed, the cells washed twice with DPBS 1X, and immunostaining was performed.

### 2.7. Mitotracker Deep Red FM staining and flow cytometry

Mitochondrial mass was evaluated using a published protocol using Mitotracker Deep Red FM (MTDR)(Mauro-Lizcano et al., 2015). Briefly, MTDR was resuspended in DMSO to obtain a stock concentration of 1mM, and stored at -20°C. On day 25 of conversion, iNs were exposed to 20µM CCCP or DMSO for 4h (Iriondo et al., 2022), media was replaced with LNM containing 250 nM of MTDR and cells were incubated for 15 min at 37°C in 5% CO_2_. Medium was then removed, cells were washed twice with DPBS 1X, and incubated for 5-10 min with Accutase at 37°C in 5% CO_2_ before being resuspended and spun down for 5 min at 400g. Supernatant was discarded, and the cell pellet was resuspended in 500µL HBSS (without calcium and magnesium) containing 1% bovine serum albumin and 0.2µg/ml DAPI (1:5000). Samples were protected from light until analysis. Samples were analyzed using a BD LSRFortessa cell analyzer. Gating was done on cells based on FSC/SSC values, doublets were removed, live cells were identified as DAPI-negative cells (BV421 filter), and mean fluorescence intensity for MTDR+ cells (APC filter) was measured.

### 2.8. Cathepsin B staining

Cathepsin B Magic Red substrate was resuspended in DMSO at a 250X stock concentration following the manufacturer’s recommendations, and stored at -20°C. Before use, the 250X stock solution was diluted to a 25X solution in sterile deionized water. iNs were incubated with a 1X working solution in LNM for 30 min at 37°C in 5% CO_2_. Cells were then washed once with DPBS 1X, and live images were acquired in LNM in a controlled environment chamber (Tokai Hit stage top incubation system, 37°C, 5% CO_2_). The density of cathepsin B was evaluated by quantifying the mean spot count of cathepsin B per neurite area, while the activity was evaluated by measuring the mean intensity of cathepsin B spots in neurites.

### 2.9. LysoFQ-GBA staining

LysoFQ-GBA was resuspended in DMSO to obtain a stock concentration of 5mM, and stored at -80°C. iNs were incubated with a working solution of 10µM of LysoFQ-GBA in LNM for 2h at 37°C in 5% CO_2_. Cells were then washed once with DPBS 1X, and live images were acquired in LNM in a controlled environment chamber (Tokai Hit stage top incubation system, 37°C, 5% CO_2_). The density of GCase was evaluated by quantifying the mean spot count of GCase per neurite area, while the activity was evaluated by measuring the mean intensity of GCase spots in neurites.

### 2.10. Pepstatin A BODIPY FL staining

Cathepsin D activity was evaluated using Pepstatin A BODIPY FL (pepA) staining. PepA was resuspended in DMSO to obtain a stock concentration of 1mM, and stored at -20°C. iNs were incubated with a working solution of 1µM of pepA in LNM for 1h at 37°C in 5% CO_2_. Medium was then removed, cells were washed twice with DPBS 1X, then fixed with 4% ice-cold paraformaldehyde (PFA) for 10 min. PFA was then removed, the cells washed twice with DPBS 1X, and immunostaining was performed. The density in LAMP1-positive structures with active cathepsin D in neurite was evaluated by quantifying the mean LAMP1-positive, cathepsin D positive colocalizing spot count per neurite area, and the activity of cathepsin D in LAMP1-positive structures was assessed by quantifying the mean intensity of the colocalizing spots.

### 2.11. Fixation and immunofluorescence staining

For cell fixation, unless otherwise mentioned, medium was removed and methanol kept at -30°C was added to the wells. The plate was transferred to -30°C for 4 min. After incubation, the methanol was removed and the cells washed twice with DPBS 1X. Cells were permeabilized and blocked for 15 min with DPBS 1X containing 5% normal donkey serum (NDS) and 0.1% Triton-X-100 before being washed twice with DPBS 1X, and incubated overnight at 4°C with appropriate primary antibodies (see **Table 2**) at proper dilution in DPBS 1X containing 5% NDS (mouse anti-ATP5a (ab14748, Abcam, 1:1000); chicken anti-MAP2 (ab5392, Abcam, 1:5000); rabbit anti-LC3B (L7543, Sigma-Aldrich, 1:500); rabbit anti-LAMP1 (L1418, Sigma-Aldrich, 1:200)). The next day, cells were washed twice with DPBS 1X, and incubated for 2h at room temperature (protected from light) with appropriate secondary antibodies diluted 1:200 in DPBS 1X containing 5% NDS and 1µg/ml DAPI (1:1000). Secondary antibody solution was removed, cells washed twice with DPBS 1x, then protected from light and kept at 4°C until image acquisition.

### 2.12. Confocal image acquisition, processing and analysis

Fluorescent images were acquired with a Nikon AX R confocal microscope. For TMRE, pepA, LysoFQ-GBA and cathepsin B, field of view images covering the whole well were acquired at 20X, then stitched to obtain a whole well image. For all other stainings, z-stack images (20 to 30 stacks, 0.2µm each, between 3 and 12 images per well) of minimum 10 iNs per cell line per condition were acquired at 60X with an oil-immersion objective. Z-stacks were deconvoluted using NIS Elements Batch Deconvolution (version 6.10.01), using the automatic settings. All analyses were done with NIS Elements Advanced Research (version 6.10.02). Neurite segmentation was done using the Segment.ai tool. Briefly, a minimum of 10 representative images of iNs per experiment were selected for each staining set. Manual tracing of neurites was done, and segment.ai was launched with 5000 iterations. After each training, visual confirmation of the accuracy of the tracing was done on random images, and training was redone with additional images in case of improper/suboptimal result. 3D reconstruction and analysis were done using the 3D tool of the software. Signal intensity threshold approach for structures of interest localized inside the neurite masks was used for object count, volume and object intensity (for pepA, LysoFQ-GBA and cathepsin B analyses), and total fluorescence intensity inside the neurite was used for TMRE analysis. For analysis normalized on the volume of neurites, volume of the 3D reconstruction of the neurite segmentation was used.

### 2.13. Statistical analyses

Statistical analyses were done using GraphPad Prism 10. Outliers were detected by Robust Regression and Outlier Removal (ROUT), with a coefficient of Q=1%. Cell lines identified as outlier values were excluded from the analysis of the parameter for which they were identified as outliers. Normality of the distribution was assessed using the Shapiro-Wilk test, and nonparametric tests were used when normality could not be assumed. Comparisons were done using either one-way ANOVA with Tukey’s multiple comparisons test or Kruskal-Wallis test with Dunn’s multiple comparisons test. To determine whether there was a significant difference between two sets of observations repeated on the same lines, a paired Student’s t-test or Wilcoxon’s test was also performed. For comparison between two groups, unpaired Student’s t-test (or Welch’s t-test in the case of significantly different variance) or Mann-Whitney’s test were used. For correlations, Pearson’s or Spearman’s correlation was used. An alpha level of 0.05 was defined for significance.

## 3. Results

### 3.1. iNs display age-related mitochondrial health impairment in neurites

Recent studies reported age-related mitochondrial health impairment in iNs generated using different reprogramming methods(Kim et al., 2018; Varghese et al., 2025). To validate the iNs generated via our reprogramming method as a model to study mitochondria-related phenotypes, we first looked at age-related mitochondrial impairment in iNs from a younger group (<60 years old) and an older group (>60 years old) of donors (**Table 1**), generated using a single reprogramming vector containing the transcription factors Ascl1 and Brn2, combined with two shRNA sequences against REST(Drouin-Ouellet et al., 2017). Because mitochondrial depolarization is one of the first events occurring following mitochondrial damage, and a reduction of the mitochondrial membrane potential (ΔΨm) is considered an indicator of mitochondrial impairment(Twig & Shirihai, 2011), we first assessed ΔΨm in our system using tetramethylrhodamine ethyl ester (TMRE), a well-known cationic lipophilic probe that accumulates specifically in polarized mitochondria(Jayaraman, 2005). While no age-associated difference in ΔΨm could be observed in the soma of iNs (**Fig. 1b**), we found a neurite-specific decrease in ΔΨm in iNs derived from older male (but not female) donors, which correlated with age (**Fig. 1c-e**). Since age-related sex differences in neurons have been reported in both human(Guebel & Torres, 2016) and animal models(Cicali et al., 2025), the remaining analyses were also disaggregated based on sex. However, unless otherwise mentioned, no further sex differences were observed.

In healthy cells, ΔΨm collapse leads to the segregation of damaged mitochondria for elimination, as well as increased mitochondrial network fragmentation, which is associated with decreased mitochondrial health(Twig & Shirihai, 2011; Zhan et al., 2013). Therefore, mitochondrial network integrity was assessed in neurites of iNs from younger and older donors, by marking mitochondria with antibodies against the ETC complex V subunit ATP5a. In line with the ΔΨm decrease observed, our results show an increase in mitochondrial network fragmentation in donors over 60 years old, as assessed by the number of ATP5a-positive puncta per MAP2-positive neurites and normalizing to neurite volume (**Fig. 1f-g**), which is in accordance with previous characterization of whole neuron mitochondrial network(Kim et al., 2018; Varghese et al., 2025). Additionally, the levels of network fragmentation were positively correlated with age (**Fig. 1h**). Together, these results support the age-related mitochondrial health impairment previously reported(Kim et al., 2018; Varghese et al., 2025).

### 3.2. iNs show age-related accumulation of mitochondria in autophagic structures in neurites following mitophagy induction

We have previously shown age-related autophagy impairment in our model(Drouin-Ouellet et al., 2022), a phenomenon further corroborated by independent studies performed on post-mortem human brain samples(Lipinski et al., 2010; Loeffler, 2019). To test whether age-related autophagy impairment would lead to the inefficient elimination of damaged mitochondria by mitophagy in iNs from older individuals, we induced mitophagy via administration of CCCP, a mitochondrial oxidative phosphorylation uncoupler that dissipates ΔΨm(Cai et al., 2012). To better understand how age affects the different stages of mitophagy, we assessed the mitochondrial load in autophagic structures at different timepoints following CCCP exposure via 3D reconstruction of immunofluorescence images. To evaluate early stages of mitophagy, we assessed translocation of mitochondria (ATP5a) into autophagosomes, using an antibody against LC3B. We found a transient increase in the proportion of LC3+ autophagosomes containing mitochondria at 2 hours following mitophagy induction in iNs derived from older individuals, as assessed by the number of ATP5a and LC3 double-positive vesicles per total number of LC3+ vesicles in neurites (**Fig. 2a-c**). This accumulation of mitochondria-containing autophagosomes was accompanied by a transient increase in the proportion of the total mitochondrial pool being engulfed in those structures 2 hours following mitophagy induction, as assessed by the colocalization of ATP5a and LC3 per total volume of ATP5a-positive staining in neurites (**Fig. 2b-c**).

Since autophagosomes fuse with lysosomes to allow for cargo degradation, we then looked at mitochondrial load in autolysosomes by measuring the colocalization of ATP5a with LAMP1. Following mitophagy induction, a persistent accumulation of mitochondria-containing LAMP1-positive compartment was seen in neurites, characterized by an increase in the density of these structures in iNs of older donors at both timepoints assessed (**Fig. 2d-h**). Additionally, in the older group, at the earlier timepoint (2 hours), the mitochondrial load increased inside the LAMP1-positive compartment, as measured by an increase of ATP5a-positive staining colocalizing with LAMP1-positive structures (**Fig. 2e-h**) and by an increase in the volume of these colocalizing fragments (**Fig. 2f-h**). At the later timepoint (4 hours), this culminated into an accumulation of LAMP1-positive compartment in neurites (**Fig. 2g-h**) (Cicali 2025,Mertens 2021). Together, these results show an accumulation of mitochondria in both autophagosomes and LAMP1-positive compartment in the neurites of iNs derived from older donors following mitophagy induction. This suggests impairment in later steps of the autophagy process, such as autolysosomal cargo degradation, leading to accumulation of mitochondria targeted for elimination.

Because the observed cargo accumulation could result in failure of mitophagy completion, we sought to measure the mitochondrial mass by flow cytometry using MTDR 4 hours following mitophagy induction. As expected, we found a significant decrease in mitochondrial mass in iNs from younger donors following mitochondrial damage with CCCP. In contrast, a great portion of lines from older donors did not show a decrease in mitochondrial mass, indicating hindered mitophagy completion (**Fig. 2i-j**). These results support age-related mitophagy impairment in neurites, characterized by an accumulation of mitochondria in autophagic structures and an incomplete clearance of these organelles.

### 3.3. Mitochondria do not accumulate in acidified autolysosomes in neurites of older donors following mitophagy induction

A recent study has shown an age-related decrease in lysosomal integrity and function in iNs(Chou et al., 2025), and age-related impairment in lysosomal acidification has often been reported in animal models(Burrinha et al., 2023; Kao, 2025). Based on these findings, we hypothesized that the incomplete clearance of damaged mitochondria in iNs derived from older donors following mitophagy induction could be due to a failure to reach functional lysosomes. Although LAMP1 is widely used for overall lysosome identification, it is not specific to functional lysosomes as it also stains endosomes(Cheng et al., 2018; Gowrishankar et al., 2015) and unacidified lysosomes(Cheng et al., 2018). Unacidified lysosomes are non-degradative, since an acidic pH is necessary for the activity of lysosomal enzymes(PILLAY et al., 2002). To specifically label acidified lysosomes, we used LDR, a dye that relies on a pH-dependent mechanism for uptake(Barral et al., 2022). We first assessed the lysosomal identity of LDR-positive structures by analyzing colocalization between LDR and LAMP1 in a village, pooling all the cell lines used for this study. As expected, all LDR-positive lysosomes were also positive for LAMP1. Interestingly, only 31.16% of LAMP1-positive structures were acidified (**Fig. 3a-b**), confirming that only a portion of LAMP1+ structures are degradative lysosomes.

To evaluate the effect of age on the translocation of mitochondria to acidified lysosomes, we assessed the colocalization of ATP5a and LDR in neurites of iNs following exposure to CCCP. The induction of mitophagy resulted in an increase in mitochondrial load inside acidified autolysosomes in neurites of the younger group, in line with the efficient completion of mitophagy observed in **Fig. 2i-j** for this age group. In contrast, no significant increase in the density of mitochondria inside LDR-positive structures in neurites from iNs of older donors was observed at any time point assessed, as measured by the volume of colocalizing ATP5a and LDR fluorescence normalized on the volume of the neurite (**Fig. 3c-d**). Because no significant difference in density and volume occupied by LDR-positive structures in neurite was found between groups at baseline or following mitophagy induction (**Fig. 3d-f**), this suggests an altered translocation of mitochondria to acidified autolysosomes in the older group, rather than an insufficient pool of acidified lysosomes. Together, these results point toward an age-associated accumulation of mitochondria in dysfunctional autolysosomes in neurites, leading to deficits in mitophagy completion.

### 3.4. Age-related mitophagy impairment is not associated with altered lysosomal enzyme activity in neurites of older donors

A study has shown that LAMP1-structures in distal neurites of cultured mice primary cortical neurons are more depleted in lysosomal proteases than their cell body counterparts, suggesting a higher heterogeneity in the degradative capacity of lysosomes in this cell compartment(Gowrishankar et al., 2015), and possibly making lysosomal structures in neurites more vulnerable to age-associated dysfunctions. This hypothesis is supported by animal studies reporting the accumulation of LAMP1-positive structures with reduced degradative capacity in the neurites of aged neurons(Burrinha et al., 2023). To investigate if the accumulation of mitochondria in LAMP1-positive and early autophagic structures following mitophagy induction could be due to decreased degradative function of lysosomes and LAMP1-positive structures, we assessed the activity of three lysosomal enzymes in neurites.

Cathepsin B is one of the most abundant lysosomal proteases (Yadati et al., 2020). In the rat brain, it is preferentially expressed in neurons (Petanceska et al., 1994). Glucocerebrosidase (GCase) is a lysosomal enzyme responsible for the lysosomal degradation of glucosylceramide (Navarro-Romero et al., 2022), and a decrease in its activity has been associated with decreased in other lysosomal enzyme function (Navarro-Romero et al., 2022) as well as autophagy defects in neurons (Schöndorf et al., 2014). Cathepsin D is a lysosomal protease whose expression, along with GCase, is increased by mitophagy induction, pointing toward its role in the degradation of damaged mitochondria (Ivankovic et al., 2016). In order to determine if an overall age-associated decrease in the degradative capacity of these enzymes in neurites could relate to the age-related mitophagy impairments observed in iNs of older donor, we assessed the density and activity of cathepsin B, cathepsin D and GCase in neurites.

No significant difference in the abundance or activity of cathepsin B (**Fig. 4a-c**) nor GCase (**Fig. 4d-f**) in neurites was found between the older and younger donors. Similarly, there was no reduction with age in the abundance of cathepsin D in LAMP1-positive structures in neurites (**Fig. 4g,i**), and no decrease was observed in the basal functionality of cathepsin D in LAMP1-positive structures (**Fig. 4h-i**). Interestingly, when disaggregating for sex, a decrease in the basal activity of cathepsin D was seen specifically in the older male donors. However, because no sex-specific difference was observed in the age-related mitophagy impairment shown in **Fig 2** and **3**, the reduced cathepsin D activity observed in older males is unlikely to account for the accumulation of mitochondria in the autophagic compartments in older donors of both sexes.

**Figure 4.**
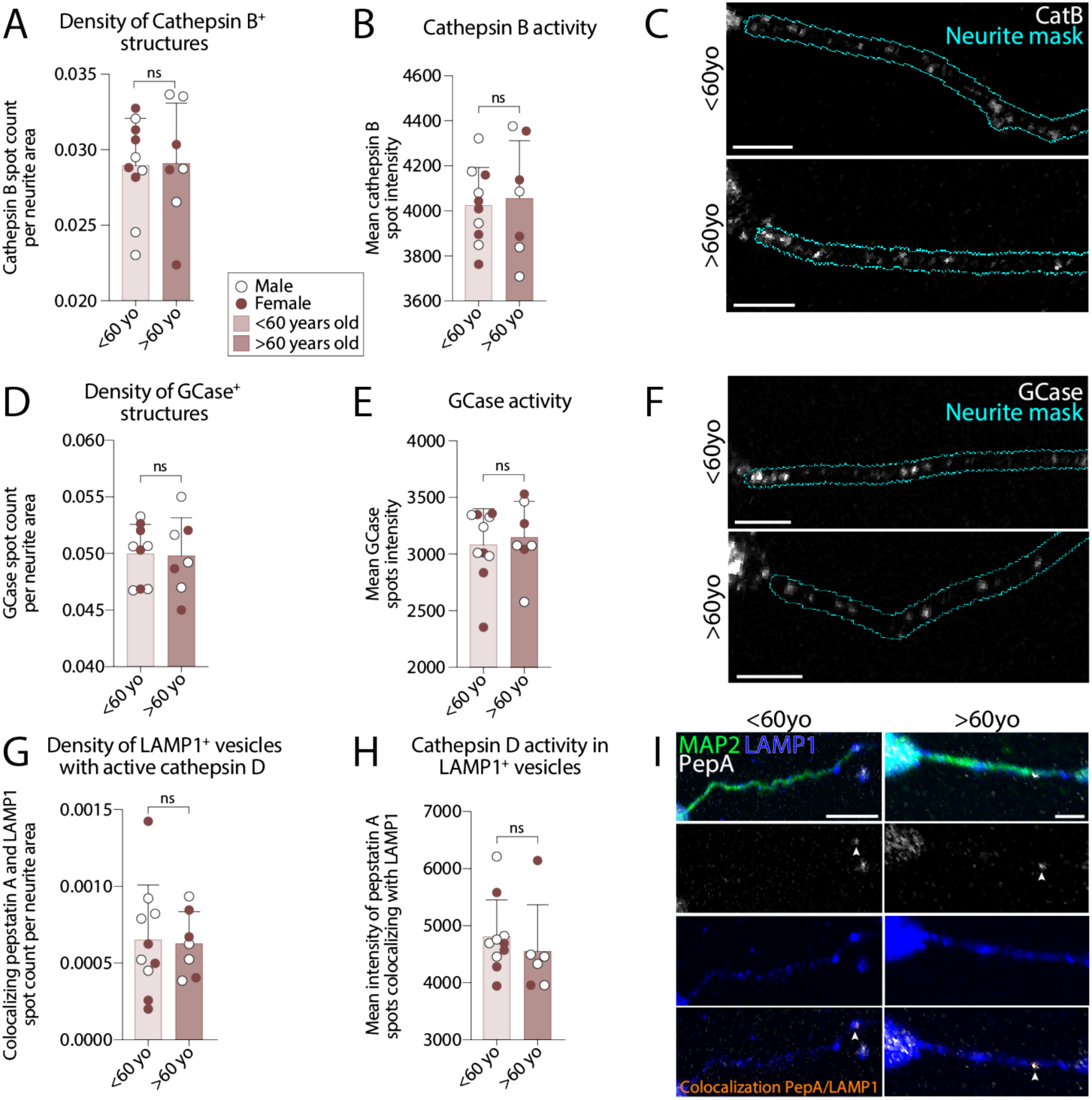
Maintenance of lysosomal enzyme function in neurites with age. **(A)** Quantification of the density (by count) of cathepsin B positive puncta per neurite area. **(B)** Quantification of the mean cathepsin B puncta intensity in neurite. **(C)** Representative fluorescence images of cathepsin B staining, and representative neurite segmentation in iNs derived from donors aged under or over 60 years old. **(D)** Quantification of the density (by count) of GCase-positive puncta per neurite area. **(E)** Quantification of the mean GCase puncta intensity in neurite. **(F)** Representative fluorescence images of GCase staining, and representative neurite segmentation in iNs derived from donors aged under or over 60 years old. **(G)** Quantification of the density (by count) of pepstatin A and LAMP1 double-positive puncta per neurite area. **(H)** Quantification of the mean pepstatin puncta intensity in pepstatin A and LAMP1 double-positive puncta in neurite. **(I)** Representative immunofluorescence images of pepstatin A and LAMP1, as well as colocalization puncta in iNs derived from donors aged under or over 60 years old. Data are expressed as mean ± the SD. Scale bars = 10µm. Abbreviations: CatB, cathepsin B; GCase, glucocerebrosidase; pepA, pepstatin A; yo, years old.

Together, these results support the hypothesis that impaired translocation of the mitochondria to acidified lysosomes, rather than impaired degradation of the mitochondria in the lysosomal compartment due to decreased lysosomal enzyme activity, would explain the accumulation of mitochondria in autophagic structures observed in the neurites of older donors. These results thus suggest that the age-associated defective mitochondrial clearance occurs upstream of lysosomal degradation rather than as a consequence of impaired lysosomal capacity.

## 4. Discussion

In this study, we hypothesized that mitochondrial health and elimination of mitochondria through the autophagy-lysosomal pathway in iNs decline in the context of normal aging. To verify this, iNs derived from young (<60 years old) and aged (>60 years old) healthy donors were generated and the ΔΨm, mitochondrial network fragmentation and lysosomal degradation of mitochondria were assessed by confocal microscopy. We report an age-related decrease in ΔΨm and an increased mitochondrial fragmentation (**Fig. 1b-h**), which is consistent with the previously reported decreased mitochondrial health with aging in human iNs and post-mortem brain samples(Bender et al., 2006; Cortopassi & Arnheim, 1990; Cottrell et al., 2001; X. Hou et al., 2018; Itoh et al., 1996; Kennedy et al., 2013; Kim et al., 2018; Kraytsberg et al., 2006; Varghese et al., 2025). Interestingly, despite being generated with a different method than that used in previous studies(Kim et al., 2018; Varghese et al., 2025), the iNs generated with our protocol display similar age-related mitochondrial phenotypes, supporting the general robustness of iNs to study mitochondria-related dysfunctions in the context of aging. We also report mitophagy impairment in the lysosomal degradation pathway of mitochondria during healthy aging, characterized by an accumulation of mitochondria in autophagosomes and insufficiently acidified LAMP1-positive compartment (**Fig. 2a-h**), culminating in ineffective mitochondrial clearance (**Fig. 2i-j**). Given that the quantity of acidified lysosomes in neurites is similar between the younger and the older group (**Fig. 3e-f**), as well as the activity of lysosomal enzymes cathepsin B, D and GCase (**Fig. 4**), this could be due to an hindered translocation of damaged mitochondria to acidified autolysosomes (**Fig. 3c-d**), or lysosomal acidification defects that have been observed in neurons in various animal models of aging(Burrinha et al., 2023; Kao, 2025). Our study provides additional support for a causal link between autophagy impairment and mitochondrial dysfunction that naturally occurs with aging in human neurons. The data presented here lay the ground for a more in-depth investigation of the underlying mechanisms of the lysosomal pathway dysregulations occurring with neuronal aging, as well as in the context of increased vulnerability of neurons in age-associated neurodegenerative diseases such as AD and PD.

The age-related mitophagy defects shown here support results from a pioneer study showing a link between aging and mitophagy in post-mortem human brain samples, which reported that the load of phosphorylated poly-ubiquitin (pSer65-Ub) structures, a marker of mitochondria targeted for degradation, increases with age(X. Hou et al., 2018), suggesting diminished mitophagy completion in older human neurons. Additionally, in iNs, a study showed a tendency towards an increase in mitochondrial mass with age(Varghese et al., 2025). Together with our study, these results point toward impaired mitochondrial clearance by the autophagy-lysosomal pathway in human neurons with aging.

In animal studies, Rubicon, Rab7 and TFEB are proteins that have been reported to be dysregulated during aging. Dysregulation of these proteins is associated with impairment of the autolysosomal degradation of cargo. Rubicon negatively regulates autophagy by blocking the fusion of autophagosomes with lysosomes through its binding with Rab7 (Bhargava et al., 2020). Increased levels of Rubicon have been reported during aging in *C. Elegans* and *Drosophila* (Nakamura et al., 2019), and its degradation, which can be induced via mitochondrial damage, increases the mitophagy flux and mitophagosome-lysosome fusion in mouse embryonic cortical neurons (Basak & Holzbaur, 2025). As such, differential basal expression of Rubicon, or differential modulation of protein levels following mitophagy induction could be linked with the accumulation of mitochondria in the insufficiently acidified LAMP1-positive compartment in iNs of older donors. Similarly, Rab7, a protein regulating the fusion of lysosomes with autophagosomes, could contribute to impaired translocation of mitochondria to acidified autolysosomes when dysregulated (Harrison et al., 2003), suggesting another possible mechanism linked to the dysfunctions observed in our study. Finally, TFEB is a master regulator of lysosomal biogenesis (Napolitano & Ballabio, 2016) and of mitochondrial quality control (S. Wang et al., 2020), and its levels have been reported to decrease with age in mice (H. Wang et al., 2021) and *C. Elegans* (Zhong & Richardson, 2025). TFEB dysregulation could underlie the age-associated impairments in mitochondrial health and mitophagy reported in this study. Further investigation of the involvement of these proteins in the regulation of lysosomal mitochondrial clearance in the neurites of human neurons during aging is warranted.

Alteration in lysosomal function during aging, as observed in animal models(Nixon, 2020), could also contribute to hindered mitophagy completion and accumulation of mitochondria targeted for degradation. Although, to our knowledge, no study in post-mortem human brain tissues has addressed changes in lysosomal function and integrity in normal aging, a recent study showed impairment in lysosomal integrity with aging in iNs(Chou et al., 2025), providing the first evidence of lysosomal dysfunction with advanced age in a human neuronal system. However, our results do not recapitulate the age-related decrease in lysosomal enzyme function reported in the literature, as we show similar density and activity of cathepsin B and D, and GCase in neurites from younger and older donors (**Fig. 4**). This discrepancy with the reported age-related phenotype might be due to the focus of this study on lysosomes specifically in neurites, which have been reported to have less degradative activity than lysosomes in the soma (Farfel-Becker et al., 2019; Gowrishankar et al., 2015). Therefore, age-associated changes in lysosomal compartment in neurites may be less pronounced than those observed in the soma, which have a higher density in lysosomes with greater degradative activity. Rather than indicating age-related impairment of lysosomal function in neurites, our results point towards altered mitochondrial trafficking to acidified autolysosomes (**Fig. 3-4**). Furthermore, we show a similar density of LDR-positive autolysosomes in neurites between the younger and older group (**Fig. 3e-f**), suggesting that lysosomal acidification is not affected by age, in line with reported data in human iNs showing an absence of lysosomal deacidification in cells derived from older donors (Chou et al., 2025). Since the proper functioning of lysosomes is not solely determined by the pH, and since numerous lysosomal enzymes other than cathepsin B and D and GCase contribute to the degradative function of lysosomes, broader investigation of the enzymatic activity of the lysosomes in which mitochondria accumulate following mitophagy induction in iNs derived from older individuals would further reinforce the hypothesis of translocation defects and rather than defects in intrinsic lysosomal function as a driver of age-related mitophagy dysfunction.

Curiously, our study shows ΔΨm impairment specifically in older males at baseline –a sex difference that has not been reported in other studies with human neurons (Kim et al., 2018; Varghese et al., 2025). Sexual hormones, such as estrogen, are known to support mitochondrial health by direct action on ETC, and by promoting the maintenance of ΔΨm via transcriptional regulation mediated by estrogen receptors α and β, amongst others(Huang et al., 2025). It has been shown that in women, levels of these receptors increase after menopause, possibly as a compensation mechanism for the decline in estrogen levels (Ishunina et al., 2007; Mosconi et al., 2024). This differential expression of estrogen receptors in females compared to males, combined with the presence of estrogen receptors agonists, such as progesterone (Burrell et al., 2026), phenol red (Berthois et al., 1986) and triiodothyronine (T3) (Nogueira & Brentani, 1996) in the neuronal medium might promote mitochondrial health in female iNs. Further investigations using a medium exempt of oestrogen/progesterone agonist, or in the presence of estrogen/progesterone antagonists such as fulvestrant and mifepristone, are warranted to decipher whether the sex difference observed is due to the *in vitro* environment or whether it is an inherent feature of mitochondrial health in neurons during aging.

Although the methods used in this study provide a broad picture of mitophagy dysfunction in the context of neuronal aging, some shortcomings remain. For instance, the use of CCCP to induce mitophagy holds several limitations, including its drastic induction of ΔΨm collapse, and its potential indirect negative effect on lysosomal function and acidification (Padman et al., 2013). However, our CCCP regimen did not lead to the collapse of lysosomal proton gradient previously observed (Padman et al., 2013), as confirmed by the uptake of LDR in acidified lysosomes following CCCP exposure in our experiments. Still, investigating mitophagy impairment using additional mitophagy inducers would strengthen our conclusions. Furthermore, although mitophagy does occur in neurites, where mitochondria are engulfed in autophagosomes before gradually fusing with lysosomes while being retrogradely transported back to the soma (Maday et al., 2012), high mitophagy activity also occurs in the soma. Notably, the somatic compartment contains most of the mitochondrial pool, and represents the site where cargo degradation is completed by autolysosomes, whether they are locally generated, or transported back from neurites (Maday & Holzbaur, 2016). Due to the compact mitochondrial network in the cell body of iNs, the analyses were carried only in neurites, where mitochondria can be thoroughly identified and segmented. An in-depth analysis of mitochondrial health and clearance in the soma would require higher resolution imaging techniques such as transmission electron microscopy, which would be low throughput and therefore challenging to do with the large number of fibroblast lines that is required to perform this type of studies on aging. A simultaneous investigation of mitophagy in the two compartments would help determine if neurite-specific processes, such as axonal transport, contributes to the observed defects. However, this phenotype is reported to occur specifically in neurites in age-related neurodegenerative diseases (Godena et al., 2014; Tammineni et al., 2017), therefore investigation in this compartment warrants specific investigation. Finally, this study focuses on the autophagy-lysosomal pathway of mitochondrial clearance. Additional studies on the impact of healthy aging on other mitochondrial quality control mechanisms, such as mitochondria-derived vesicles and mitochondrial transfer, would allow for a more complete understanding of age-associated phenotypes regarding the elimination of damaged mitochondria in neurons.

Together, our results show age-related mitophagy deficiency in the neurites of iNs derived from individual older than 60 years old, which is linked to autolysosomal dysfunction. Further investigation of the mechanism underlying the defective progression and completion of mitophagy observed in our model may reveal novel targets anti-aging therapies, as well as for the treatment of age-associated neurodegenerative disorders.

## 5. Statements

### 5.1. Ethic statement

aHDFs (see **Table 1**) were obtained from Coriell Institute and used under full local ethical approval: CERSES-18-004-D (University of Montreal).

### 5.3. Author contributions

EL: conceptualization, methodology, formal analysis, investigation, visualization, writing –original draft, writing –review and editing. JPIG: investigation, visualization, writing –review and editing. JDO: conceptualization, methodology, visualization, writing –review and editing.

### 5.5. Acknowledgments

We thank Nicolas Giguère (University of Montreal) for his help with confocal imaging, and Chiara Tocco (University of Montreal) for manuscript revision.

## 7. Supplementary materials

**Supplementary Figure 1.**
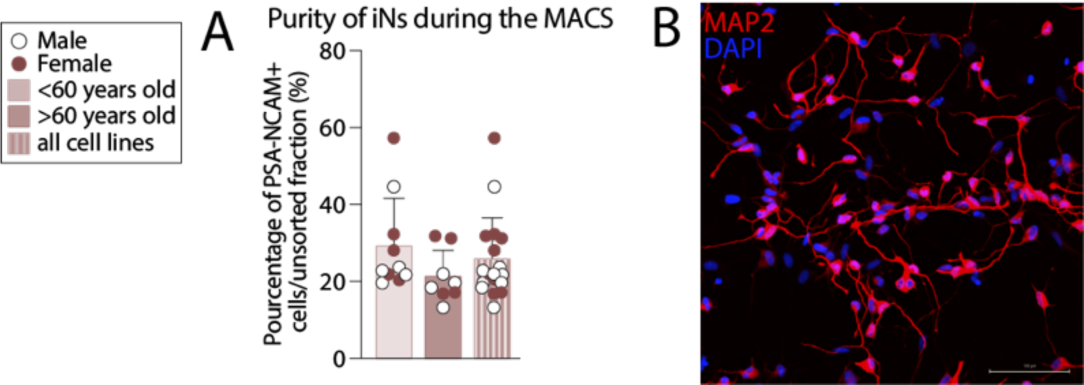
Neuronal purity of the cultures before and following purification procedure by MACS. **(A)** Quantification of the number of cells in the PSA-NCAM-positive fraction of the MACS on the total cell count before sorting. **(B)** Representative confocal imaging of DAPI-positive and MAP2-positive cells.

